# Cross validation coordinate meta-analysis: contrast analysis

**DOI:** 10.1101/2024.12.16.628634

**Authors:** C.R. Tench

## Abstract

Coordinate based meta-analysis (CBMA) can be used to estimate where a future neuroimaging study testing a particular hypothesis might report summary results (activation foci, for example). However, current methods cannot be validated for all possible analyses because of empirical features that might not be appropriate. Furthermore, the use of voxel-wise null hypothesis significance testing (NHST) in the algorithms is not in keeping with meta-analysis, where statistical significance is secondary to the primary aim of effect estimation.

Cross-validation coordinate analysis (CVCA) has been described, which can eliminate the need for the empirical use of spatial uncertainty and avoid voxel-wise p values. The result is an estimated effect that is not based on voxel-wise statistical significance, and which allows the uncertainty to reduce with larger numbers of studies as expected.

Here an additional function of CVCA is detailed, which uses cross-validation to contrast two different meta-analyses produced using two sets of studies (*A & B*) to identify differences. Such contrast analysis is common in CBMA. The results are a contrast image with regions that differentiate studies *A* from studies *B*.

Software to perform CVCA is freely available.

## Introduction

### What is CBMA

Coordinate based meta-analysis (CBMA) is commonly used to summarise effects reported by multiple independent, but related by a shared hypothesis, neuroimaging studies. It is employed to meta-analyse (amongst others) voxel-based morphometry (VBM) or functional magnetic resonance imaging (fMRI) and uses only reported summary results such as coordinates (foci of activation, for example). By analysing multiple studies simultaneously, those results that represent a common finding among at least some of the studies are identified and are assumed indictive of relevant and reliably measurable effects. In the absence of whole brain statistical images with which to perform full image based meta-analysis (IBMA), CBMA can provide a useful summary of where studies testing the same hypothesis are likely to report significant results.

Popular CBMA methods that use only coordinates are the activation likelihood estimation (ALE) method (1,2), and the kernel density estimation (KDE) methods (3). Other methods utilise the effect estimate (Z score or *t* statistic) reported for each coordinate and use a null hypothesis of zero effect (4,5). These, however, have stricter inclusion criteria than those requiring only coordinates in that the studies must report effects uncensored by effect sign; they must include both activation and deactivation for example. Indeed, the common aim of meta-analysing only activations (or only deactivations) would not be possible, and neither would mixing studies that test only for activation with studies that test for both activation and deactivation; these could violate the null hypothesis even in the absence of a meta-effect. A further class of CBMA algorithms involve Bayesian methods (6,7), but these involve specification of priors by experts.

In meta-analysis the aim is to obtain better estimates of an effect that has been measured independently multiple times. Effects typically have a size and variance, and the meta-analysis uses both. In CBMA the effects are spatial, but there is no accompanying spatial variance. This leaves multiple independent effects per study without any obvious way to relate those reported in one study to those reported in other studies. Therefore, in CBMA an empirical estimate of unknown spatial variance is introduced. This, along with null hypothesis significance testing (NHST), is used to cluster the independent findings into an estimated meta-effect. Unfortunately, fixing the empirical spatial uncertainty does not account for the inherent uncertainty of the studies, nor does it consider the need to modify that uncertainty of the estimated effect for the number of studies; many studies should lead to more certainty than few studies. An alternative method (cross validation coordinate analysis; CVCA) uses cross-validation to automatically estimate a spatial uncertainty instead of using an empirical value (8). This estimate may then reduce as the number of studies increase, as expected. CVCA also eliminates the need for voxel-wise p-values, null hypothesis significance testing (NHST), and subsequent type 1 error control, which are secondary aims in meta-analysis where estimation is the primary goal.

A typical application for CBMA is contrast analysis (9), where studies of type *A* and studies of type *B* are compared to produce a spatial estimate of difference. Like CBMA this is typically conducted using fixed empirical uncertainty estimates and voxel-wise p-value thresholding. Here a cross-validation approach to contrast analysis is described as an extension to CVCA. Software to perform CVCA is freely available https://github.com/TenchCR/NeuRoi/blob/main/neuroi.zip

## Methods

CVCA uses a kernel density estimation method to describe the density of reported foci from a set of studies. The kernel is based on a beta distribution parameterised to be Gaussian-like, but with limited extent preventing distal effects interacting. The details of CVCA have been described previously (8). But briefly, in CVCA the kernel is parameterised as having a width (*ω*). For a given width meta-analysis is conducted on N-1 studies and a score, which reflects how closely foci from the remaining held-out study agree with the meta-results, is computed. This score is averaged over different held-out studies, and the kernel width that maximises it is chosen. This imposes the condition that the meta-results should reflect the held-out data, which is a major difference to other CBMA algorithms that look for statistical significance. Consequently, instead of becoming nonsignificant due to too few studies, the meta-effect uncertainty is increased as expected for a meta-analysis.

For contrast analysis the kernel width for studies of types *A & B* are first estimated independently using CVCA. Using this estimate an image is created per study, where the image intensity is defined by the kernel due to the nearest focus in the study; this is analogous to the modelled activation image used in the ALE algorithm (10). These images are summed and divided by the number of studies to produce a likelihood-of-reported effect image. For two sets of studies being analysed (*A & B*) these images are subtracted to indicate where foci from studies of type *A* (*B*) are more likely than foci from studies of type *B* (*A*). The task is then to estimate a threshold on this image that would distinguish *A* from *B*. This is achieved using pairs of leave out studies (one of type *A* and one of type *B*) to define a threshold that maximises a score reflecting the distinction between study types on average.

### Definitions

Studies are of type *A* or *B*. The kernel widths for each study type are *ω*_A_ and *ω*_B_ respectively and are established by CVCA. For study *i* of type *A* an image is formed

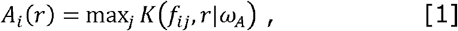

where *r* is the location in the image, *f*_ij_ is the position of the *j*^th^ focus in study *i*, and *K* is the kernel parameterised by width *ω*_A_. Note that only foci identified as above threshold in the CVCA algorithm are included in this image. Equivalent images are computed for studies of type *B*.

A likelihood image for each type of study is computed by summing over these

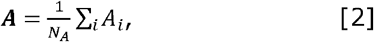

where *N*_A_ is the number of studies of type *A*, which standardises the image for contrast analysis. A similar image is computed for studies of type *B*.

The difference between study types is represented by a difference image

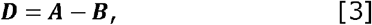

which is positive where studies of type *A* are more likely to report effects, and negative where studies of type *B* are more likely to report.

### Thresholding

The difference image **D** represents the difference between studies of types *A & B*. It requires a threshold to localise regions that can distinguish these. Given the threshold the contrast image is

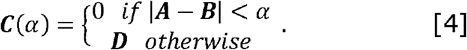

Where this image is positive (negative) it indicates greater likelihood of reported effect in studies of type *A* (*B*). To compute the threshold *α* cross validation is used. For each held-out pair of studies, the value of the image **C**, given the threshold, at each focus in the held-out studies is computed. For the held-out study of type *A* (*B*), a focus falling where **C** is positive increases a count *A*_+_ (*B*_-_) by 1, while a count *A*_-_ (*B*_+_) is incremented by 1 if **C** is negative. The score for each held-out pair is the proportion of foci falling in the respective part of image **C**(*α*),

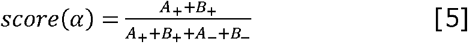

and this is averaged over all held-out pairs. This score summarises the proportion of foci, of those that fall in none zero regions of **C**, that fall in the part most likely given the type of study reporting. If studies of type *A & B* report similarly, the score is expected to be around 0.5. Where there is strong distinction between study type, the score will tend towards 1.

### Statistical significance

The results are subjected to a single significance test, producing only one p-value and eliminating the need for any multiple test correction. Firstly, the score is computed for the original study sets *A & B*. Then, the studies are permuted to form sets *A*’ and *B*’, consisting of the same number of studies as *A & B*, and the score recomputed; a sample of a null score. Multiple samples of the null score are computed and used to calculate a Z score for the contrast between studies *A* and *B*, which can subsequently be converted to a single p-value. A significant result would typically have a Z score of around 2 or more.

### Organising studies for analysis

When conducting CBMA the foci from multiple studies must be extracted. Within study it is common that multiple experiments are performed on the same subjects, often with similar hypotheses. These are not independent and cannot be considered so in the meta-analysis without causing bias. A proposed solution is to arrange all foci by study, rather than experiment (11). However, a consequence for methods that spatially randomise foci independently to generate the null distribution, such as the popular ALE algorithm, is a reduction in the sensitivity to meta-effects due to the number of foci randomised being larger than the actual number of independent effects (12). The solution suggested here is to include only one experiment from each study. The other experiments could then be used to perform sensitivity analysis to make sure results are robust to perturbation.

### Experiments

Two experiments are conducted for demonstration purposes, and to show how results are presented and interpreted. Data are extracted from previously published work developing a minimum Bayes threshold for CBMA (13). Firstly, two random groups of VBM Schizophrenia studies are extracted and compared; it is expected that there is no difference between these. For the second experiment fMRI studies of time perception are compared to fMRI studies of face discrimination, which is expected to reveal differences. The original data are modified to include only one experiment from each study.

For assessment of statistical significance 20 analyses of permuted studies are conducted. For an accurate estimate of Z, ideally more iterations would be performed. However, the score itself is also directly interpretable and possibly more important than Z, therefore, 20 is generally sufficient; this is a user selected option, and the analysis takes only a few minutes to conduct so large numbers of permutations are computationally feasible if wanted.

## Results

### Schizophrenia

The Schizophrenia data available from (13) consists of 49 VBM studies. These were split randomly into groups of 21 studies and 28 studies. Figure 1 depicts example results from CVCA on all 49 studies, and figure 2 shows where the split groups differ according to contrast analysis. It would appear that there are differences between these groups despite their equivalence. However, foci from the held-out studies do not preferentially fall in regions to indicate real difference, hence the score (equation 5) is 0.52, and not statistically significant (Z=0.76).

**Figure 1.**
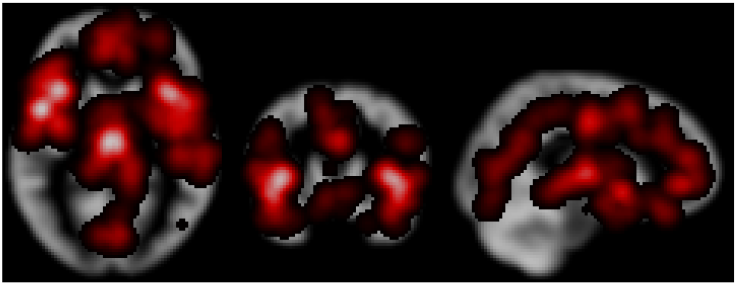
The result, shown for example orthogonal slices, of cross validated coordinate analysis on a set of 49 VBM studies of Schizophrenia.

**Figure 2.**
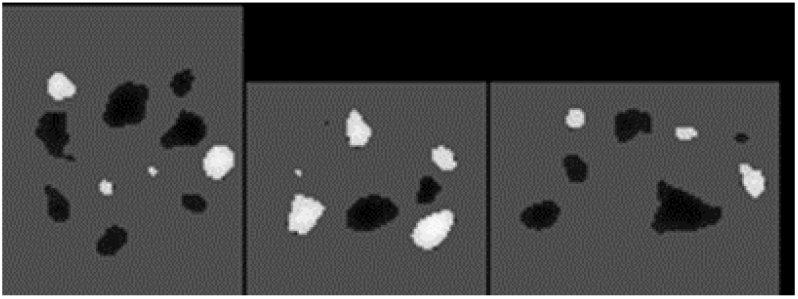
The Schizophrenia studies were split randomly into two groups of 21 and 28 studies. The figure shows the result of contrasting these groups, which as expected was not significant giving a score of 0.52 (Z=0.76).

### Time perception v face discrimination

The face discrimination group includes 18 fMRI studies (CVCA results depicted in figure 3), and the time perception group includes 85 fMRI studies (CVCA results depicted in figure 4). The contrast between these two study groups reveals regions that can distinguish the groups (figure 5). Analysis results show that on average studies of type A will generally report foci in the positive regions while studies of type B will generally report foci in the negative regions.

**Figure 3.**
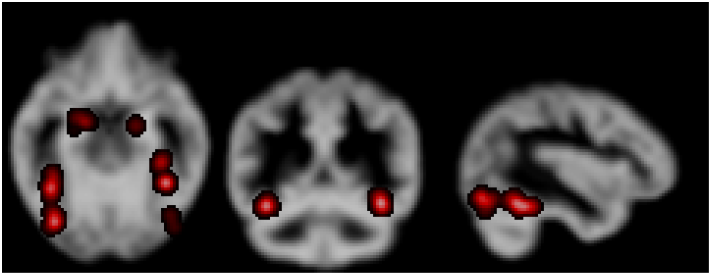
CVCA results on 18 fMRI studies of face discrimination.

**Figure 4.**
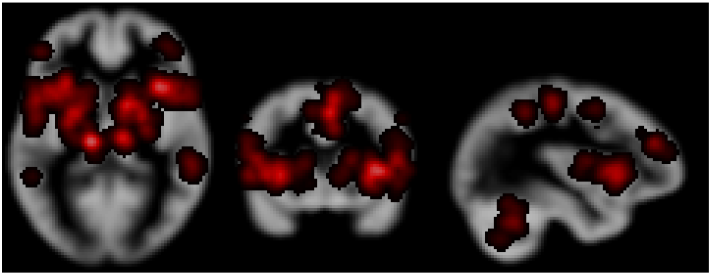
CVCA results on 85 fMRI studies of time perception.

**Figure 5.**
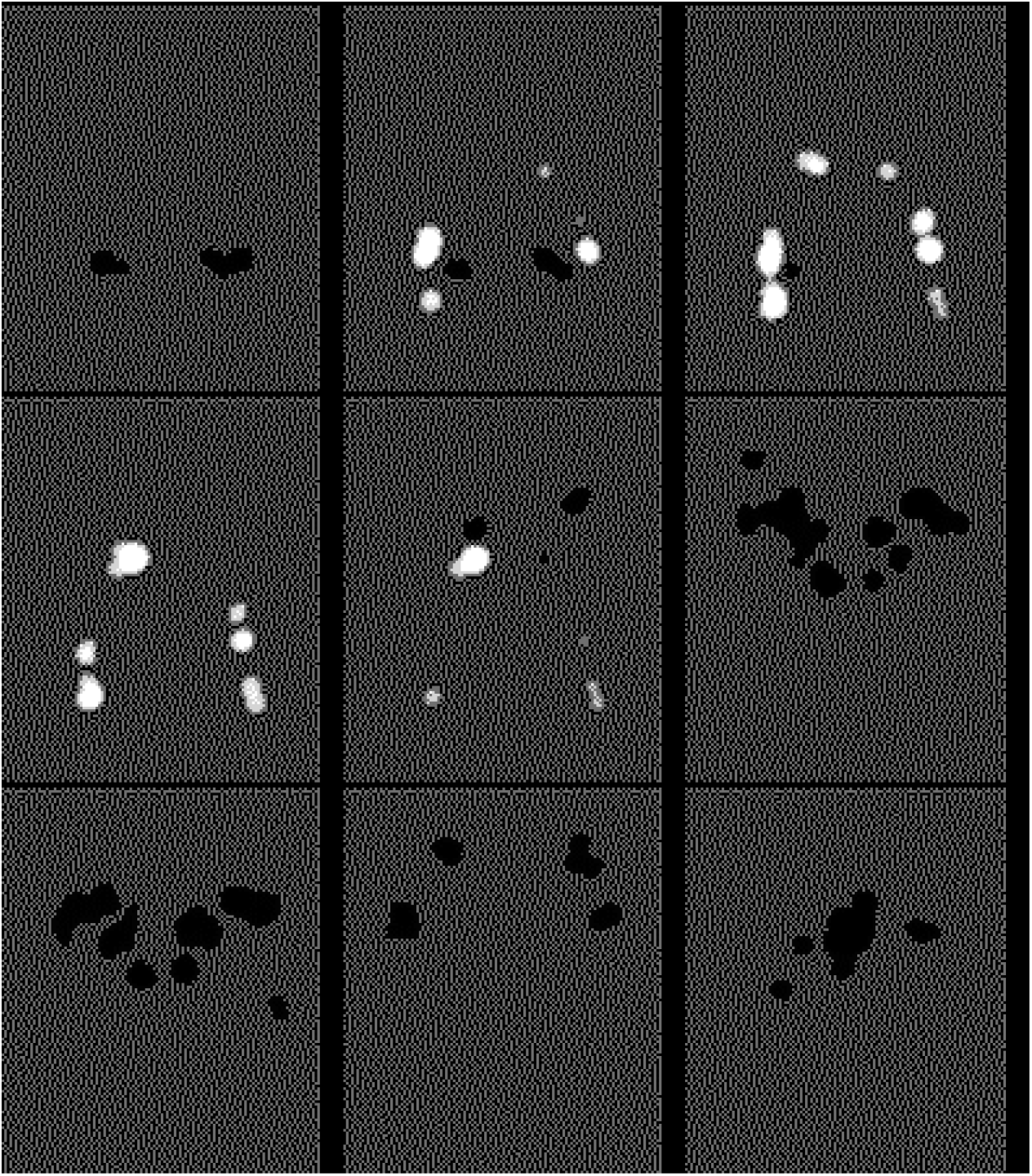
Selected image slices from the results of contrasting face discrimination with time perception. The positive (white) regions indicate where studies of face discrimination often report, but studies of time perception generally don’t. The negative (black) regions indicate where studies of face discrimination rarely report, but studies of time perception often do. The maximum score (equation 5) for this difference is 0.82, indicating strongly contrasting effects. This contrast was statistically significant with Z=5.34.

## Discussion

Here a CBMA contrast algorithm has been presented that does not rely on null hypothesis significance testing to threshold spatial results and does not use an empirically fixed kernel width that might not be valid for all cases. Its aim it to extend cross validated coordinate analysis to contrast two study groups and produce an anatomical map of regions that distinguish the groups. A cross-validation approach is used to avoid voxel-wise p-values and subsequent thresholds, which are not in keeping with meta-analysis but are typical in CBMA algorithms.

The use of empirical kernel widths in CBMA algorithms means that results may not be valid under some conditions. For example, if the meta-analysis involves very many independent studies, it is likely that a smaller kernel would be appropriate (14) to indicate how the reported foci in different studies are related. Conversely, a meta-analysis involving few studies would likely require a larger kernel as the mean distance to the nearest foci from the different studies would be larger. Statistical thresholding using NHST is normal practice in CBMA, and in neuroimaging in general, and is a pragmatic solution to a problem. However, such thresholding is most interpretable in a prospective setting, where an experiment can be designed to force (with some small type 2 error rate) a correctly hypothesised effect to be significant. In meta-analysis however, there is no way to modify the sample of data analysed to define the power of the study.

Results presented here demonstrate how contrast analysis using CVCA works on reported imaging study results. When studies testing the same hypothesis are compared no difference is found, reflected in a small Z score. However, when studies of different hypotheses are compared, a contrast image detailing where the differences are in the brain is produced with a significant Z score of 5.34. A useful property of the method is a score that indicates how different the two groups are in a number ranging from about 0.5 (no difference) to 1(completely distinct). By removing the baseline 0.5 from this score, a distance matrix contrasting multiple study groups could be formed and principle coordinate analysis performed to place all groups relative to each other in a low dimensional space.

The issues of meta-analysing neuroimaging results relate to lack of spatial variance for the reported effect. Without this variance, there continues to be limitations compared to conventional meta-analysis. However, a kernel density approach is a classic way of estimating the spatial density, and cross-validation a classic method of parameterising the kernel properties to the data. This is the approach taken by CVCA to eliminate an empirical variance estimate. Furthermore, meta-analysis is not typically a problem of statistical significance, but rather one of estimation. CVCA eliminates the voxel-wise p-values in favour of a maximisation of an interpretable function, using cross-validation, to produce the results. In the case of contrast analysis discussed here cross-validation is used to optimise a function that would distinguish studies of 2 different hypotheses. The score being optimised is quantitatively interpretable in that a score of around 0.5 means there is little to distinguish the reported results, while a score approaching 1 means the reports from the different test of the hypotheses can readily be distinguished.

## Summary

Conventional meta-analysis uses a reported variance that has no equivalent in CBMA. Typically, an empirical value is substituted, but this is a strong limitation. CVCA estimates a variance that allows effects in different studies to be combined subject to a sensible constraint of best reflecting held out data. In the case of contrast analysis the aim is to distinguish studies of two different hypotheses. The use of cross-validation in CVCA and its extension to contrast analysis is an attempt to eliminate two limitations of other CBMA algorithms: the use of an empirical kernel that might not be valid for all analyses, and voxel-wise p-values, which are not typically an aim of meta-analysis.

